# Reduction of DNA Topoisomerase Top2 reprograms the epigenetic landscape and extends health and life span across species

**DOI:** 10.1101/2024.11.14.623693

**Authors:** Man Zhu, Meng Ma, Lunan Luo, Feiyang Li, Jiashun Zhang, Yan Pan, Lu Yang, Ying Xiao, Ziyan Wang, Bo Xian, Yi Zheng, Hao Li, Jing Yang

## Abstract

DNA topoisomerases are essential molecular machines that manage DNA topology in the cell and play important roles in DNA replication and transcription. We find that knocking down the enzyme Topoisomerase Top2 or its mammalian homolog Top2b increases the life span of *S. cerevisiae*, *C. elegans*, and mice. Top2b reduction also extends the health span of mice and alleviates the pathologies of aging in multiple tissues. At the cellular/molecular level, Top2b reduction attenuates the major hallmarks of aging, such as cellular senescence, de-regulated nutrient sensing, epigenetic alteration, and lysosomal biogenesis. We observed that Top2b reduction significantly changes the epigenetic landscape in various mouse tissues toward those of the young animals, and differentially down-regulates genes with active promoter and high expression. Our observations suggest that Top2 reduction confers longevity effect across species via a conserved mechanism, and may be used as a novel therapeutic strategy for countering aging.

## Introduction

The proportion of the global population aged 65 years and above is projected to increase from 10 percent in 2022 to 16 percent by 2050 (*1*). The rapid increase in the elderly population has drawn worldwide interest in efforts to combat aging. Over the past few decades, the application of molecular genetics to model organisms has led to the discovery of several biological pathways that regulate life span across species, leading to insight into the mechanisms of aging and the development of potential therapeutic interventions (*2, 3*). An important revelation from these studies is that some of the longevity pathways have essential cellular functions and are important for growth and development; however, their down-regulation post-development significantly extends life span. Examples include the classic insulin/IGF signaling and the TOR pathways; their discovery led to the development of promising potential anti-aging drugs currently under human clinical trial (*2*).

To further our understanding of the mechanisms of aging and to identify alternative therapeutic targets, it is desirable to explore other key molecular machines/pathways for their role in regulating life span across species. However, information on the life span phenotype of essential genes is scarce, especially for mammalian species. In the simple model organism yeast, the life span of the non-essential gene knock out mutants has been measured systematically through a multi-year effort and ∼200 mutants with extended life spans were identified (*4, 5*). As a significant fraction of the non-essential gene knock out mutants have been profiled transcriptionally (*5*), we analyzed the correlation between the gene expression changes and the life span of the mutants and identified a number of essential genes whose down-regulation strongly correlates with extended life span across multiple mutants (**Supplementary Table 1**). Among the top hits is a DNA topoisomerase Top2, with an essential function in managing DNA topology and regulating replication and transcription. Combined with the previous observation that reducing Top2 extends life span in yeast (*6*), we hypothesized that Top2 serves as a key node in the gene regulatory network that influences life span and that reducing Top2 might extend life span across species through a conserved mechanism.

Yeast Top2 has two mammalian homologs, TOP2A and TOP2B. While TOP2A is primarily expressed in proliferating cells and is crucial for DNA replication, TOP2B is expressed in all cell types and plays a more prominent role in chromatin remodeling and transcriptional regulation that is closely tied to aging. We thus decided to focus on TOP2B. TOP2B is an essential double-stranded DNA topoisomerase, pivotal in identifying DNA topological configurations and relieving DNA torsional strain via cutting, rotating, and reconnecting DNA strands (*7*). Emerging research underscores its importance in chromosome architecture maintenance, DNA replication and repair, and transcriptional regulation (*7, 8*). TOP2B has been much less studied in the context of aging, with a few previous studies implicating it in age-related retinopathies and hearing impairment (*9, 10*), and down-stream response to dietary restriction (*11*).

In this study, we investigate whether reduction of Top2 or TOP2B confers longevity phenotype across species and explore the potential mechanisms. We found that knocking down Top2 or TOP2B extends the life span of yeast, *C. elegans*, and mice. TOP2B reduction also extends the health span of mice, and alleviates the characteristics and pathologies of aging in multiple tissues. At the cellular/molecular level, Top2 or TOP2B reduction attenuates the major hallmarks of aging, such as cellular senescence, deregulated nutrient-sensing, epigenetic alterations, and lysosomal biogenesis. We observed that TOP2B reduction significantly alters the epigenetic landscape in various mouse tissues toward those of the young animals, and differentially down-regulates genes with active promoter and high expression. Our observations suggest that Top2 or TOP2B reduction confers longevity effect via remodeling of epigenetic and transcriptional landscapes and suppression of aberrantly expressed genes in old cells. Our findings also suggest TOP2B can be a novel therapeutic target for countering aging.

## Results

### Reduction of Top2 or its vertebrate homolog Top2b extends the life span of yeast, worms, and mice

We first thoroughly elucidated the evolutionary relationships of Top2 homologous genes by conducting an unrooted phylogenetic tree constructed by the neighbor-joining method within MEGA 11 software across 14 distinct species. All discerned Top2 proteins were categorized into four distinct groups, highlighting the conserved evolution of both Top2 and Top2b (**Extended Data Fig. 1A**).

To investigate whether TOP2 knock down extends life span across species, we first test whether we can replicate the previously observed life span extension by Top2 knock down in yeast (*6*). We constructed Top2 DAmP (Decreased Abundance by mRNA Perturbation) strains in both the BY4741 (Mat-a) and BY4742 (Mat-alpha) genetic backgrounds. The DAmP strains have decreased abundance of the target mRNA due to 3’ modification that renders it less stable (*12*). Compared to the wild-type (WT) strains, the Top2 mRNA was knocked down by 41% and 44% in the TOP2 DAmP strains with BY4741 and BY4742 backgrounds respectively (**Fig. 1A and 1B**). Compared to the BY4741 (mean RLS=23.92, n=164) and BY4742 WT (mean RLS=24.98, n=110) strains, the mean RLS of top2 DAmP strains in BY4741 (mean RLS=28.10, n=166) and BY4742 (mean RLS=30.28, n=104) backgrounds was extended by 17.5% and 21.2%, respectively (**Fig. 1C**, all *P*<0.0001), as measured by using the traditional microdissection technique.

**Figure 1.**
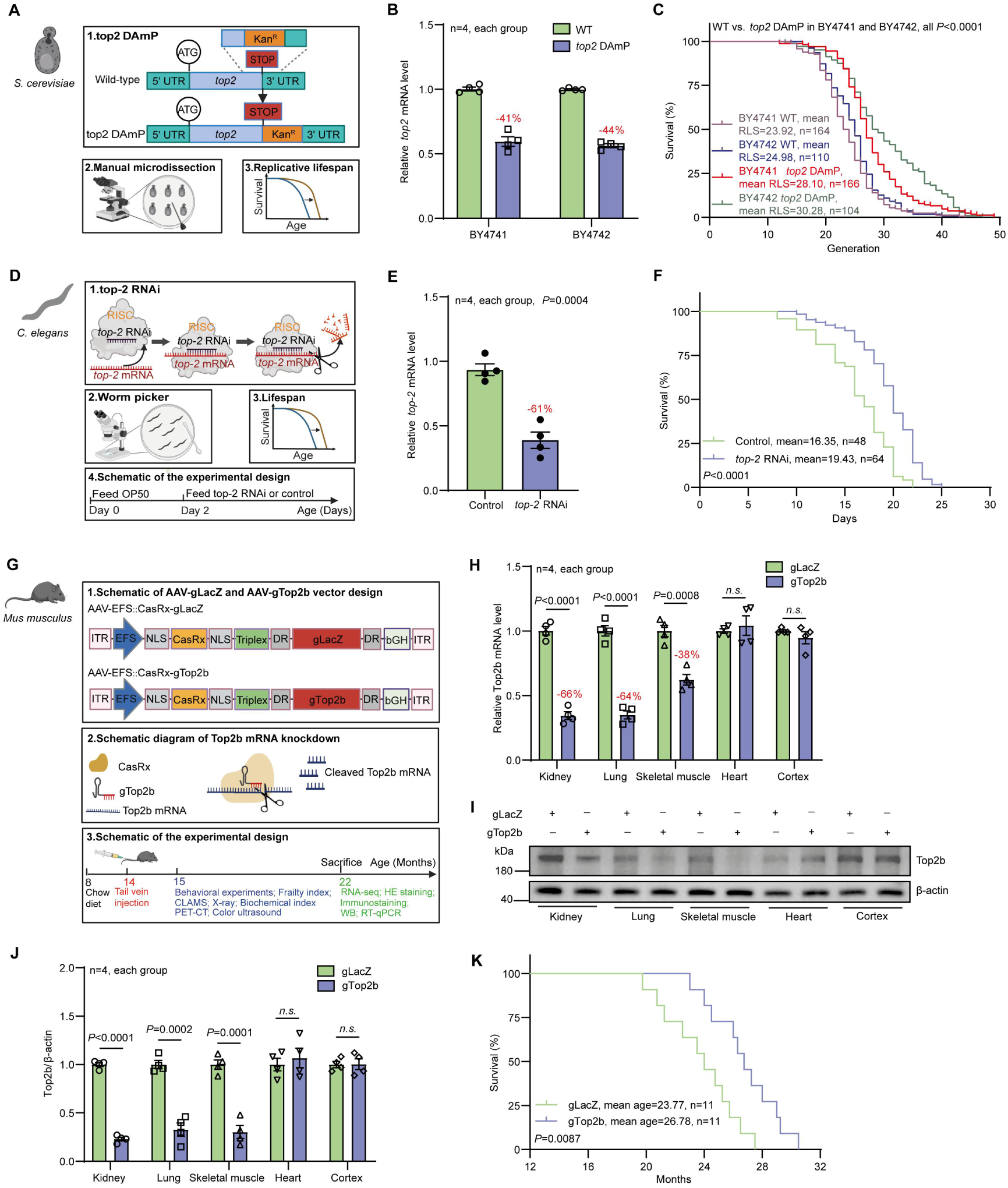
Reduction of Top2/Top2b extends the life span of yeast, worm, and mice. (A) Schematic of *top2* DAmP approach in *S.cerevisiae*. DAmP, decreased abundance by mRNA perturbation; Kan^R^, kanamycin resistance. (B) Relative *top2* mRNA levels in BY4741 and BY4742 wild type (WT) strains and the corresponding *top2* DAmP mutants, as measured by RT-qPCR. (C) Replicative life span (RLS) of BY4741 and BY4742 WT strains and the top2 DAmP mutants. (D) Schematic of *top-2* RNAi approach in *C. elegans*. RNAi, RNA interference; RISC, RNA-induced silencing complex. (E) Relative *top-2* mRNA levels in *C. elegans* for the control group and the *top-2* RNAi groups, as measured by RT-qPCR. (F) Life span of the *C. elegans* for the control and *top-2* RNAi groups. (G) Schematic of Top2b knockdown by the CRISPR/CasRx system in mice. AAV, adeno-associated virus. (H) Relative Top2b mRNA levels in the kidneys, lungs, skeletal muscles, hearts, and cortices of mice in the gLacZ control and gTop2b groups, measured by RT-qPCR. (I-J) Relative Top2b protein levels in the kidneys, lungs, skeletal muscles, hearts, and cortices of mice in the gLacZ control and gTop2b groups, as measured by western blot. (K) The life span of C57BL/6 mice for the gLacZ control and gTop2b groups. Statistical analysis was performed using GraphPad Prism v8.0 software (https://www.graphpad.com). Data were considered statistically significant at *P* < 0.05 calculated by using Student’s t-test (B, E, H, and J) or log-rank test (C, F, and K). *n.s.* indicates not significant. All values are means ± SEM. The corresponding n values (number of cells, worms, or mice) are shown within each sub-plot.

We next determined the effect of Top2 reduction on the life span of *C. elegans* using the N2 strain. We employed an RNAi approach (**Fig. 1D**) to knock down the Top2 mRNA. With the RNAi approach, it is possible to analyze dose-response of the life span vs. the degrees of knock down by feeding worms with different proportions of *E. coli* with the RNAi construct and the wildtype *E. coli* (HT115 strain). We experimented with a range of different proportions of *E. coli* with TOP2 RNAi (0%, 25%, 50%, 75%, 100%) and found life span extension in a dose-dependent manner, with 50% yielding the largest life span extension (**Extended Data Fig. 1B and 1C**). Compared to the control (mean life span=16.35 days, n=48), the life span of the 50% Top2 RNAi group (mean life span=19.43 days, n=64) was extended by 18.8% (**Fig. 1E and 1F**, *P*<0.0001); the mRNA expression level of Top2 was reduced to 61% of the WT level (Fig. 1E). Interestingly, the optimal dose for the life span also yielded the best health indexes such as the number of pharyngeal beat per minute and percent of worms entered the slow swallowing phase (**Extended Data Fig. 1D-1F**). We therefore focused on 50% Top2 RNAi in the subsequent worm experiments.

Top2 is evolutionarily conserved in multicellular eukaryotes. Top2a and Top2b are two distinct forms in vertebrate species. Top2a is primarily detected in actively dividing cells such as germ cells and is crucial for DNA replication. In contrast, Top2b is widely and highly expressed in various tissues and cell types (*13*) and plays an important role in regulating chromatin structure and gene expression. To investigate whether TOP2 knock down extends life span in mammals, we decided to focus on TOP2B and analyzed the effect on life span by TOP2b knock down in mice. We designed the CRISPR/CasRx system to knock down Top2b and constructed AAV DJ serotype viruses to deliver the TOP2b knock down vector and the LacZ control into multiple organs/tissues through tail vein injection at 14 months age of the mice (**Fig. 1G**). Real-time quantitative PCR (RT-qPCR) and Western blot analysis revealed that compared to the control group (gLacZ), the levels of Top2b mRNA (**Fig. 1H**) and protein (**Fig. 1I and 1J**) in the kidneys, lungs, and skeletal muscles of Top2b knockdown mice (gTop2b group) were downregulated, despite no significant impact in the cortex and heart. In comparison to the gLacZ group (mean age=23.77 months, n=11), the average life span of gTop2b group mice (mean age=26.78 months, n=11) was extended by 12.7% (**Fig. 1K**, *P*=0.0087).

### Top2b/Top2 knockdown improved the health span of mice and *C. elegans*

An effective anti-aging strategy should both extend life span and improve health span. The mouse frailty index (FI) is a comprehensive compilation of health indicators, encompassing body weight, coat condition, grip strength, mobility, vision, and hearing; lower FI indicates a more healthy condition (*14*). To assess the impact of Top2b knockdown on the health span of mice, we conducted FI scoring every two months and performed extensive behavioral assays to evaluate the health status of the mice.

No discernible differences in FI scores were observed between the two groups before AAV virus injection (at 14 months of age), however, FI scores of mice in the gTop2b group were significantly lower than those in the gLacZ group after intravenous injection (**Fig. 2A**, all *P*<0.01). At 20 months of age, a hair assessment was performed. We observed that mice in the gTop2b group displayed a reduced incidence of alopecia/depigmentation phenotype and less pronounced spinal kyphosis compared to the gLacZ group (**Fig. 2B and 2C**), and it seemed that Top2b knockdown had a limited effect on mice body weight (**Fig. 2D)**.

**Figure 2.**
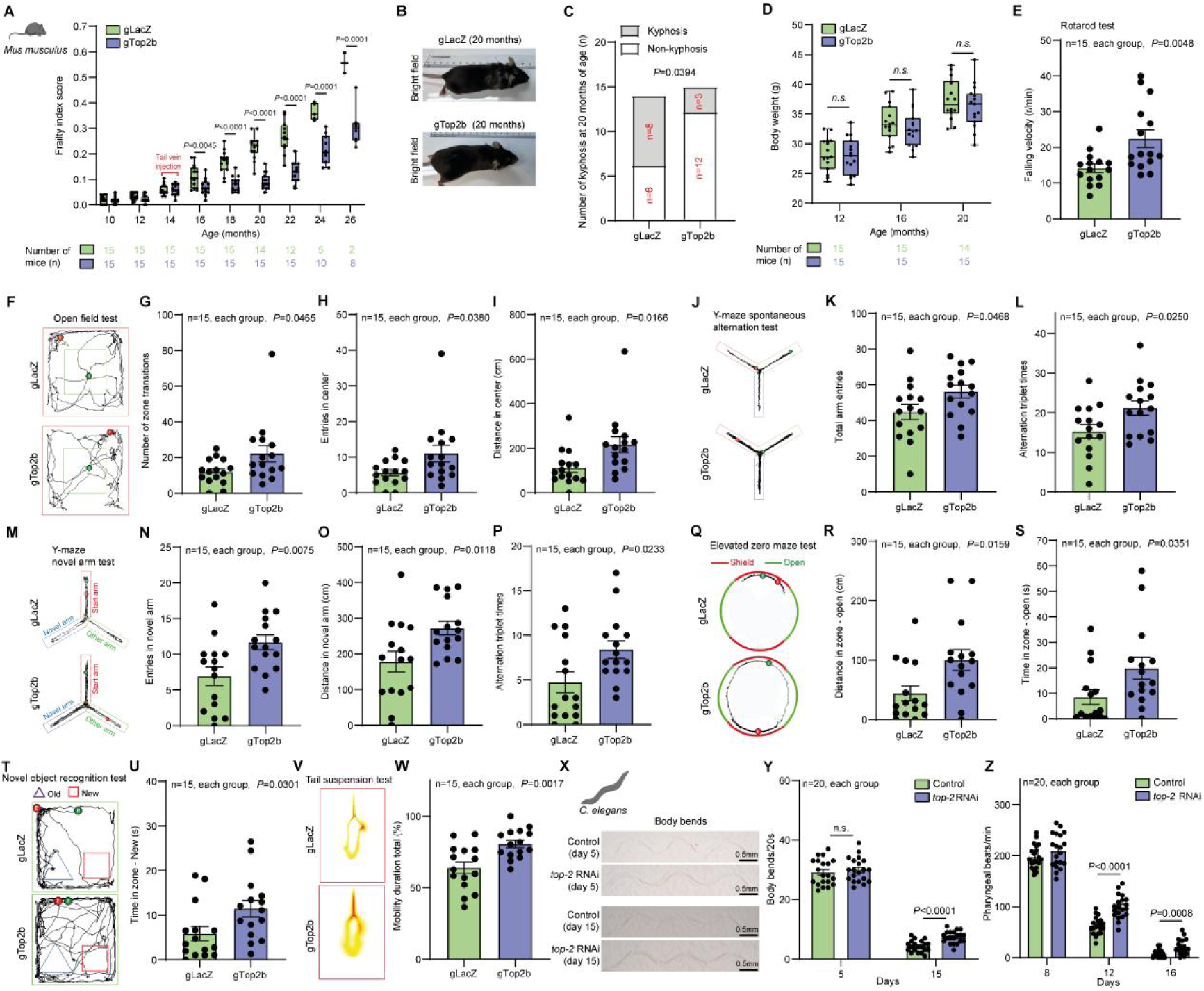
Top2b/Top2 reduction improves the health span of mice and worms. (A) Frailty index (FI) scores for C57BL/6 mice in the gLacZ and gTop2b groups were recorded at 10 to 26 months of age, with assessments conducted once every two months. The number of mice assessed at each age is listed at the bottom of the plot. (B) Bright-field images of gLacZ and gTop2b mice. (C) Number of kyphosis in C57BL/6 mice of the gLacZ and gTop2b groups at 20 months of age. (D) Body weight of C57BL/6 mice in the gLacZ and gTop2b groups at 12, 16, and 20 months of age. (E) Falling velocity in rotarod tests for C57BL/6 mice in the gLacZ and gTop2b groups. (F-I) Representative images of open field tests and number of zone transitions, entries in the center, and distance in the center for C57BL/6 mice in the gLacZ and gTop2b groups. (J-L) Representative images of Y-maze spontaneous alternation tests and total arm entries and alternation triplet times for C57BL/6 mice in the gLacZ and gTop2b groups. (M-P) Representative images of Y-maze novel arm tests and entries in novel arm, distance in novel arm, and alternation triplet times for C57BL/6 mice in the gLacZ and gTop2b groups. (Q-S) Representative images of elevated zero maze tests and distance in zone-open and time in zone-open for C57BL/6 mice in the gLacZ and gTop2b groups. (T-U) Representative images of novel object recognition tests and time in zone-new for C57BL/6 mice in the gLacZ and gTop2b groups. (V-W) Representative images of tail suspension tests and mobility for C57BL/6 mice in the gLacZ and gTop2b groups. (X-Z) Representative images of body bend and bending frequency and pharyngeal pumping rate in *C. elegans* fed with control RNAi and top-2 RNAi after 5 and 15 days. Statistical analysis was performed using GraphPad Prism v8.0 software (https://www.graphpad.com). Data were considered statistically significant at *P* < 0.05 calculated by using Student’s t-test (A, D, E, G-I, K, L, N-P, R, S, U, W, Y, and Z) or Chi-squared test (C). *n.s.* indicates not significant. All values are means ± SEM or n (%). The corresponding n values (number of mice or worms) are shown in each sub-plot.

Muscle coordination, endurance, and strength decline with age (*15*). In the rotarod test, mice in the gTop2b group showed superior performance at both latencies to fall and falling velocity compared to the gLacZ group (**Fig. 2E, Extended Data Fig. 2A**). Reduced memory and exploration and increased anxiety are common behavioral changes induced by aging (*16*). We conducted extensive behavioral experiments including the open field test (**Fig. 2F-I, Extended Data Fig. 2B-D**), Y-maze spontaneous alternation test (**Fig. 2J-L, Extended Data Fig. 2E and 2F**), Y-maze novel arm test (**Fig. 2M-P, Extended Data Fig. 2G**), elevated zero maze test (**Fig. 2Q-S, Extended Data Fig. 2H-J**), and novel object recognition test (**Fig. 2T and 2U, Extended Data Fig. 2K**). We found that compared to the gLacZ group, mice in the gTop2b group demonstrated significantly enhanced spatial exploration and memory, and exhibited lesser anxiety. Although depressive-like behavior is not a typical manifestation of aging, aging undoubtedly serves as a crucial precipitant (*17*). The tail suspension test also revealed that mice in the gTop2b group exhibited less depressive-like behavior compared to the gLacZ group (**Fig. 2V and 2W**).

We also evaluated the benefit of Top2 knockdown to the health span of *C. elegans*. Bending frequency and pharyngeal pumping rate represent two pivotal metrics for evaluating age-associated physiological deterioration in *C. elegans*, both exhibiting a progressive decline with advancing age. We found a significant increase in both bending frequency and pharyngeal pumping rate of *C. elegans* in the Top2 RNAi group compared to the HT115 group (**Fig. 2X-Z**).

### TOP2B reduction mitigates the characteristics and pathologies of aging in multiple tissues in mice

To analyze the effect of top2b knock down in different tissue/organs, we evaluated the morphology of the kidneys, lungs, livers, hearts, skins, and skeletal muscles of gTop2b mice using hematoxylin-eosin (HE) staining. Compared to the gLacZ group (23-month-old) mice, gTop2b (23-month-old) mice exhibited alleviation of glomerular atrophy in kidney (**Fig. 3A and 3B**), more regular hepatic lobule structure and reduced anisokaryosis in liver (**Fig. 3E and 3F**), and decreased average thickness of alveolar septa in lung (**Fig. 3G and 3H**). Additionally, in gTop2b mice, there was an enlargement of the cross-sectional area of skeletal muscle fibers (**Fig. 3C and 3D**), an increase in the number of nuclei in the left ventricular wall (**Fig. 3K and 3L**), and thickening of the dermis and epidermis layers and increased number of hair follicles in skin (**Fig. 3I-J**).

**Figure 3.**
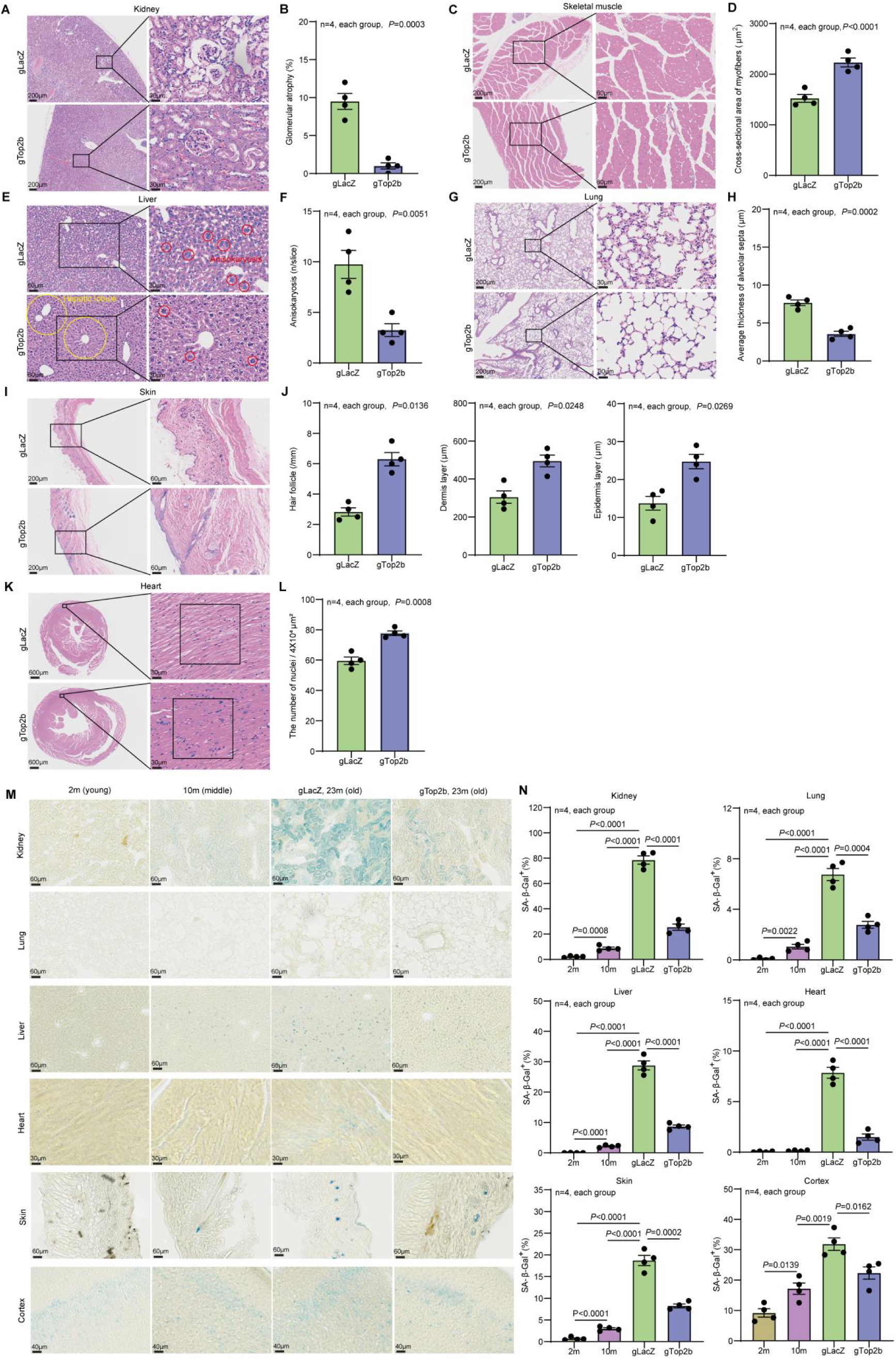
Top2b reduction mitigates the characteristics and pathologies associated with aging in multiple mouse tissues. Histology of various tissues of 23-month-old mice in the gLacZ control group and gTop2 group is analyzed (A-L). (A) The longitudinal sections of the kidney tissues of were stained with HE. The right panels show a magnified view of the boxed area in the left panels. Scale bars represent 200 μm and 30 μm for the left and right panels. (B) Average glomerular atrophy. (C) The transverse sections of mouse skeletal muscle were stained with HE. The right panels show a magnified view of the boxed area in the left panels. Scale bars represent 200 μm for the left panels and 60 μm for the right panels. (D) Average cross-sectional area of myofibers. (E) Mouse liver tissues were stained with HE. The right panels show a magnified view of the boxed area in the left panels. Scale bars represent 60 μm for the left panels and 30 μm for the right panels. The yellow circles highlight the hepatic lobule. The red circles indicate anisokaryosis. (F) Average anisokaryosis. (G) Mouse lung tissues were stained with HE. The right panels show a magnified view of the boxed area in the left panels. Scale bars represent 200 μm for the left panels and 30 μm for the right panels. (H) Average thickness of alveolar septa. (I) Mouse skin tissues were stained with HE. The right panels show a magnified view of the boxed area in the left panels. Scale bars represent 200 μm for the left panels and 60 μm for the right panels. (J) Average hair follicle density, average dermis layer thickness, and average epidermis layer thickness in mouse skin. (K) The short axes of cardiac tissues from mice were stained with HE in the left ventricular wall regions. The right panels show a magnified view of the squared area in the left panels. Scale bars represent 600 μm and 30 μm for the left and right panels. (L) The number of nuclei in each region of the left ventricular wall. (M) Representative images of SA-β-Gal staining of various tissues from young (2m old), middle age (10m old), old (23m old) gLacZ control, and old (23m old) gTop2b groups. (N) quantitation of percent of SA-β-Gal positive cells in various tissues. Statistical analysis was performed using GraphPad Prism v8.0 software (https://www.graphpad.com). Data were considered statistically significant at *P* < 0.05 calculated by using Student’s t-test (B, D, F, H, J, and L) or one-way ANOVA (N). All values are means ± SEM. The corresponding n values (number of mice) are shown within each sub-plot.

Furthermore, we performed SA-β-Gal staining and identified SA-β-Gal positive cells in kidneys, lungs, livers, hearts, skins, and cortex of 2-month-old wild-type mice, 10-month-old wild-type mice, 23-month-old gLacZ, and 23-month-old gTop2b mice **(Fig. 3M and 3N)**. The analysis revealed an age-dependent increase in the percentage of SA-β-Gal positive cells. Administration of gTop2b treatment via tail vein injection resulted in a significant reduction in the percent of SA-β-Gal positive cells in all these tissues compared to the gLacZ mice (**Fig. 3M and 3N**). These data indicate that Top2b knockdown effectively reduces senescent cells and alleviates age-associated characteristics in all the tissues we analyzed. It is worth noting that, although the gTop2b AAV injection via the tail vein didn’t decrease the Top2b protein level in the heart and cortex (**Fig. 1I and 1J**), the percent of SA-β-Gal positive cells still declined in these two tissues relative to the control, suggesting that targeting Top2b in some of the tissues can produce a systemic effect that is communicated to other tissues.

### TOP2B knockdown alleviates various cellular aging hallmarks

To examine the effect of Top2b knowdown at the cellular/molecular level, we analyzed various aging hallmarks. A prominent hallmark of aging is cellular senescence -- the permanent proliferation arrest mediated by activation of cyclin-dependent kinase inhibitors (CKIs) CDKN1A/p21 and CDKN2A/p16 (*18*). We thus analyzed the protein expression of p21 and p16 in replicative-, stress- or oncogene-induced-senescence of human IMR-90 cells. Compared with young cells (the 6^th^ generation), p16 and p21 are up-regulated in older cells (the 12^th^ generation) and by stress- or oncogene-induced-senescence (**Fig. 4A and 4B**). Etoposide-induced senescence exhibited a stronger p21 increase, while oncogenic K-RAS^G12V^-induced senescence showed a more significant p16 increase, indicating a distinctive response to different senescence-inducing signals. We observed that both p16 and p21 were attenuated to a much lower level when Top2b was knocked down (**Fig. 4A and 4B**).

**Figure 4.**
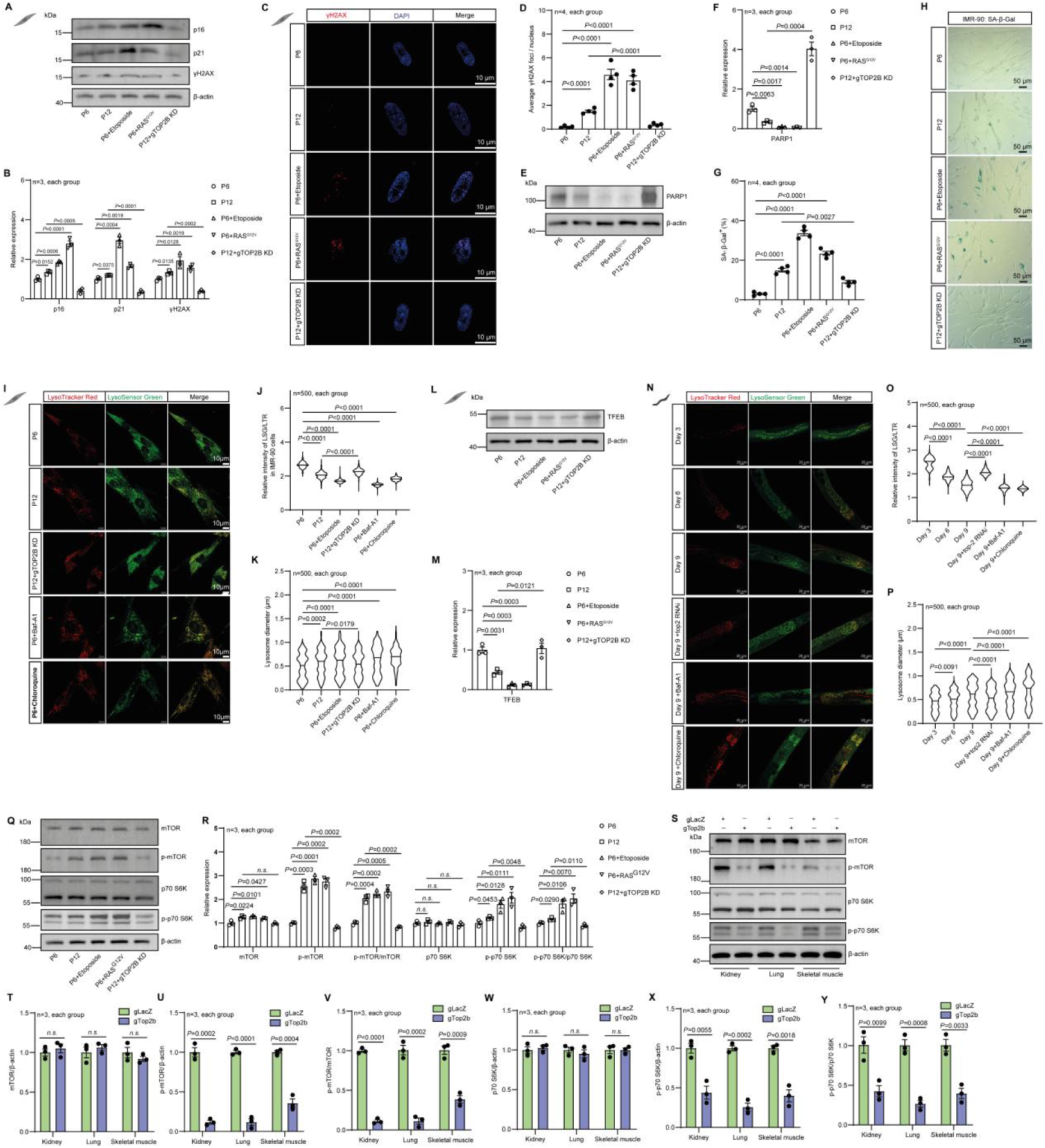
Top2b knock down reduces various cellular aging hallmarks in human IMR-90 cells and in various mouse tissues. (A-H) IMR-90 cells were induced to undergo replicative, stress-induced, or oncogene-induced senescence. The protein levels of p16, p21, and γH2AX were detected by western blotting (A, B). γH2AX foci in DAPI-stained nuclei of IMR-90 cells were measured by immunofluorescent staining (C, D). The protein level of PARP1 was detected by western blotting (E, F). SA-β-Gal staining (blue-stained cells) and quantification of percent of SA-β-Gal positive cells were shown in G, H. (I-P) Confocal fluorescence images of IMR-90 cells (I-K) and *C. elegans* (N-P) stained by LysoTracker Red DND-99 (LTR) and LysoSensorTM Green DND-189 (LSG), and the fluorescence intensity ratio of LSG/LTR was measured as an indicator of lysosomal acidity. (L, M) IMR-90 cells were induced to undergo replicative, stress-induced, or oncogene-induced senescence. The protein level of TFEB was detected by western blotting. The relative intensity of LSG/LTR and average lysosome diameter in IMR-90 cells (J, K) and *C. elegans* (O, P) were quantified. (Q-R) Western blot analysis of nutrient-sensing mTOR signaling proteins in IMR-90 cells. (S-Y) Western blot analysis of nutrient-sensing mTOR signaling proteins in mouse kidney, lung, and skeletal muscle. Statistical analysis was performed using GraphPad Prism v8.0 software (https://www.graphpad.com). Data were considered statistically significant at *P* < 0.05 calculated by using Student’s t-test (T-Y) or one-way ANOVA (B, D, F, G, M, R) or Kruskal-Wallis test (J, K, O, P). All values are means ± SEM. The corresponding number of samples for IMR-90 cells, number of worms, and number of mice are shown within each sub-plot.

DNA damage accumulation is another cellular aging hallmark. DNA double-strand breaks (DSBs) are potent inducer of cellular senescence, triggering a cascade of DNA damage response pathways that lead to irreversible growth arrest and the acquisition of senescent phenotypes (*19*). We found that the number of serine-139-phosphorylated H2AX (γH2AX) foci, a marker of DSB, significantly increased in replicative-senescent, oncogenic K-RAS^G12V^, and etoposide-induced senescent IMR-90 cells and was markedly attenuated after Top2b knockdown (**Fig. 4C and 4D**). The total amount of γH2AX as measured by western blotting showed the same trend (**Fig. 4A and 4B**). Poly(ADP-ribose) polymerase 1 (PARP1) is the central enzyme for Poly-ADP-ribosyl production which is protective to cells during DNA damage (*20*). We found that PARP1 protein decreased in replicative-senescent, oncogenic K-RAS^G12V^, and etoposide-induced senescent IMR-90 cells, but significantly increased after Top2b knockdown (**Fig. 4E and 4F**).

We observed increased expression of SA-β-Gal activity in replicative-senescent (P12), oncogenic K-RAS^G12V^, and etoposide-induced senescent P6 IMR-90 cells relative to the P6 IMR-90 control cells. Top2b knockdown in P12 cells significantly reduces SA-β-Gal compared to P12 control cells (**Fig. 4G and 4H**). SA-β-Gal is generally linked to the increased content of lysosomes (*21*), and the concomitant decline of lysosomal function correlates with cellular senescence. To analyze whether knockdown of Top2b can improve aging-related lysosome dysfunction, we examined lysosome diameter and function in replicative-senescent and Top2b knockdown IMR-90 cells by co-staining with LysoTracker Red DND-99 (LTR) and LysoSensorTM Green DND-189 (LSG), and using Chloroquine (a lysosome inhibitor) and Baf-A1 (a V-ATPase inhibitor) treatments as positive controls. The fluorescence intensity ratio of LSG/LTR was measured as a positive indicator of lysosomal acidity. Replicative aging, as well as Chloroquine or Baf-A1 intervention, resulted in an increase in lysosome diameter and a decrease in LSG/LTR ratio, while Top2b knockdown in P12 cells led to a decrease of lysosome diameter and an increase of LSG/LTR ratio relative to the P12 control cells (**Fig. 4I-K**).

TFEB is a master regulator of lysosome biogenesis and overexpression of TFEB can induce lysosomal exocytosis (*22*). Inhibition of mTORC1 can reduce phosphorylation of several serine residues of TFEB permitting its nuclear translocation and transcriptional function (*23*). We found that in replicative-senescent, oncogenic K-RAS^G12V^, and etoposide-induced senescent IMR-90 cells, TFEB protein was decreased. Top2b knockdown significantly increased TFEB level in P12 cells **(Fig. 4L and 4M)**, further indicating improved lysosome function with Top2 reduction. To examine whether the effect of Top2b knockdown on lysosome is conserved, we analyzed the effect of Top2 RNAi on lysosome diameter and function in *C. elegans*. The aging process (worms of 3,6 and 9 days) and Chloroquine or Baf-A1 treatments led to an increase in lysosome diameter and a decrease in LSG/LTR ratio, while Top2 RNAi had an opposite effect, ameliorating aging-related lysosome dysfunction (**Fig. 4N-P**). In summary, the above results indicate that, as an important aging hallmark, lysosomal function decreases with age, including an increase in diameter and a decrease in acidity, while the knockdown of Top2b or Top2 can significantly reverse these trends both in human IMR-90 cells and in worms.

Deregulated nutrient sensing is another major hallmark of aging. Aged cells exhibit a deregulated capacity to sense nutrients and poor metabolic status, while effective nutrient sensing contributes to delaying senescence (*24*). Given the crucial role of mTORC signaling in longevity-related metabolism regulation and nutrient-sensing, we quantified the protein level of p-mTOR and mTOR, and its downstream indicator P70-S6K in IMR-90 cells. We found that p-mTOR/mTOR and p-P70-S6K/P70-S6K ratios increased in replicative-senescent, oncogenic K-RAS^G12V^, and etoposide-induced senescent IMR-90 cells relative to the P6 control cells, while Top2b knockdown in P12 cells significantly reduced the two ratios back to those in P6 cells (**Fig. 4Q and 4R**), indicating that Top2b knockdown significantly reduces mTORC signaling.

We further tested whether Top2b knockdown reduces mTORC signaling *in vivo* in mice. Top2b knockdown mice exhibited significantly decreased p-mTOR/mTOR ratio and p-P70-S6K/P70-S6K ratio in the kidney, lung, and skeletal muscle of gTop2b mice compared to the gLacZ mice, although the mTOR level and the P70-S6K level themselves did not change, indicating decreased mTORC signaling (**Fig. 4S-Y**).

Together, our data suggest that Top2b knockdown significantly reduces mTORC pathway signaling both in human cells and in mice. It is known that decreasing mTORC signaling through genetic manipulations or through drug inhibition (such as Rapamycin) is an effective life span-extending intervention.

### TOP2 down-regulation contributes to global transcriptional changes indicative of longevity promotion

To explore the potential molecular mechanism underlying the longevity effect of Top2/Top2b knockdown, we performed RNA-seq analyses of Top2 RNAi-treated *C. elegans* (**Fig. 5A-D; Supplementary Table 2**) and various tissues of Top2b knockdown mice including kidney, lung, and skeletal muscle (**Fig. 5E-O, Supplementary Table 3-5**).

**Figure 5.**
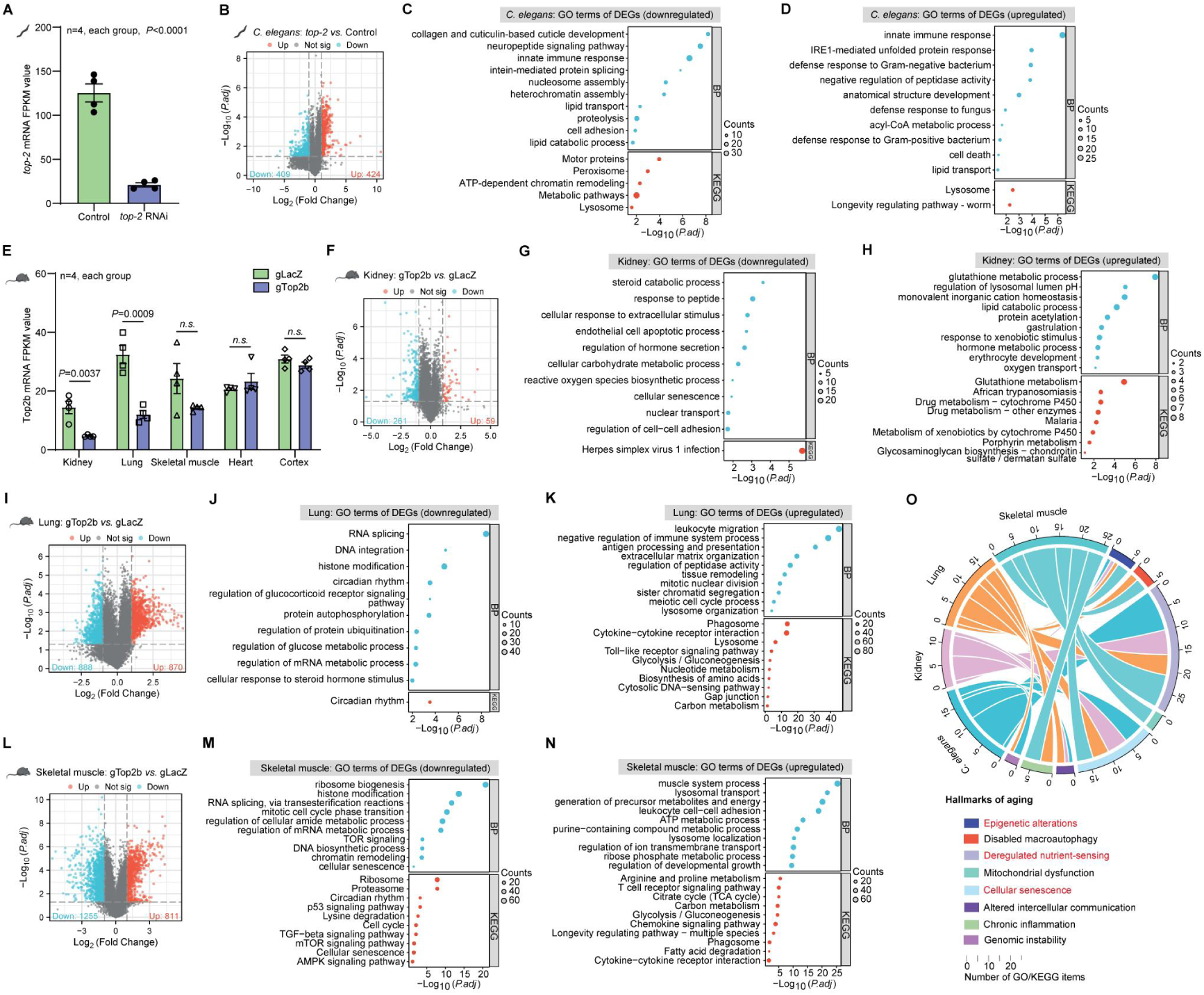
Top2b knockdown led to changes of the global transcriptional program targeting multiple aging hallmarks. (A) The adjusted Top2 FPKM levels between the control and *top-2* RNAi-treated *C. elegans*. FPKM, fragments per kilobase of exon per million mapped fragments. (B) Transcriptome analysis of the upregulated (up) and downregulated (down) DEGs between control and *top-2* RNAi-treated *C. elegans*. (C, D) GO analysis identified up (C) and down (D) regulated functional categories in DEGs between control and *top-2* RNAi-treated *C. elegans*. BP, biological process; KEGG, Kyoto Encyclopedia of Genes and Genomes. (E) The adjusted Top2b FPKM levels in the kidney, lung, and skeletal muscle from gLacZ and gTop2b mice (F) Transcriptome analysis of the up and down DEGs in the kidney tissues between gLacZ and gTop2b mice. (G, H) GO analysis identified up (G) and down (H) regulated categories in the kidney tissues between gLacZ and gTop2b mice. (I) Transcriptome analysis of the up and down DEGs in the lung tissues between gLacZ and gTop2b mice. (J, K) GO analysis identified up (J) and down (K) regulated categories in the lung tissues between gLacZ and gTop2b mice. (L) Transcriptome analysis of the up and down DEGs in the skeletal muscle tissues between gLacZ and gTop2b mice. (M, N) GO analysis identified up (M) and down (N) regulated categories in the skeletal muscle tissues between gLacZ and gTop2b mice. (O) Circos plot illustrating connections between DEGs due to TOP2b knockdown in different tissues and hallmarks of aging. Statistical analysis was performed using GraphPad Prism v8.0 software (https://www.graphpad.com). Data were considered statistically significant at *P* < 0.05 calculated by using Student’s t-test (A and E) or Fisher’s exact test and adjusted by the Benjamini-Hochberg method (C, D, G, H, J, K, M, and N). Genes with |log_2_fold change (FC)| > 1 and adjusted *P* value (by the Benjamini–Hochberg method) < 0.05 were considered DEGs (B, F, I, and L). *n.s.* indicates not significant. All values are means ± SEM. The corresponding n values (number of mice) are shown within each sub-plot.

We observed global changes of transcription profiles in the Top2/Top2b knockdown mutants compared to the wild-type control. In Top2 RNAi-treated *C. elegans*, Top2 mRNA abundance was reduced to 20% of the WT level (**Fig. 5A**), and a total of 424/409 genes were upregulated/downregulated (|Log_2_FC|>1; p value<0.05, **Fig. 5B, Supplementary Table 6**). Genes up-regulated are significantly enriched for several biological processes (BP), including innate immune response and IRE1-mediated unfolded protein response, indicating increased immune defense and stress resistance. On the other hand, genes down-regulated are significantly enriched for collagen and cuticulin-based cuticle development, indicating a down-regulation of developmental/growth-related process (**Fig. 5C and 5D**).

CRISPR-mediated Top2b knockdown via tail vein injection of AAV virus selectively decreased Top2b mRNA in the lung, skeletal muscle, and kidney, but not in the heart and cortex, based on the high throughput sequencing data **(Fig. 5E)**. This is consistent with the RT-qPCR and protein expression results (**Fig. 1H-J**). Compared with the gLacZ control mice, gTop2b mice have 59/261, 870/888, and 811/1255 genes upregulated/downregulated in the kidney, lung, and skeletal muscle respectively (|Log_2_FC|>1; *p* value<0.05, **Fig. 5F, I, L, and Supplementary Table 7-9**).

Functional enrichment analyses indicate that the GO BP or KEGG terms enriched in differentially expressed genes (DEGs) differ in different tissues. However, the general theme is up-regulating processes important for the specific tissue function and down-regulating protein synthesis/general growth. For example, “glutathione metabolic process” is up-regulated in kidney, where glutathione plays an important role in anti-oxidant and de-toxification, which is a key function of kidney **(Fig. 5G and 5H).** In the lung, the most highly enriched categories in the up-regulated genes include “leukocyte migration”, “negative regulation of immunce process”, and “anti-gene processing and presentation”, centered around immune response to infectious agents or irritants. The most significantly down-regulated category in lung is “RNA splicing”, a key step in protein synthesis (**Fig. 5J and 5K**). Skeletal muscle has the biggest change of transcriptome based on the number of DEGs. “ribosomal genesis” is the most highly enriched in the down-regulated genes, indicating down-regulation of protein synthesis. Genes responsible for muscle function and energy production are up-regulated (**Fig. 5M and 5N, Extended Data Fig. 3, Extended Data Fig. 4**).

We further compared the DEGs across different mouse tissues. For up-regulated DEGs, there is almost no overlap among the three tissues (**Extended Data Fig. 5A**). In kidney, a group of genes related to N-acetyltransferase activity including Nat8, Nat8f5, nat8f6, and nat8f2 were the most significantly up-regulated. This is a family of enzymes whose primary role is in the formation of mercapturic acids and in detoxification pathways in liver and kidney (*25*). The most up-regulated genes in lung were related to immune response. Examples include Acod1, responsible for catalyzing the production of itaconate, an immunoregulatory metabolite, and several cytokines (Cxcl2, Ccl4, Ccl3, Cxcl10). In skeletal muscle, the most significantly increased genes are related to muscle function. Examples include Actc1, which encodes cardiac muscle alpha-actin, and Casq1, which serves as the primary calcium-binding protein in the sarcoplasmic reticulum of skeletal muscle. Interestingly, spermidine biosynthesis-related genes Amd1 and Smox are highly induced, and spermidine is a known longevity molecule that promotes autophagy (*26*).

Interestingly, there are 23 shared genes in the down-regulated DEGs across tissues (shown in the Venn Diagram, **Extended Data Fig. 5B-F)**. Among them are several zinc finger transcription factors whose function are not well characterized, including zfp932, zfp518a, and zfp52. Other genes include Resf1, which positively regulates DNA methylation-dependent heterochromatin assembly, Dek, which is involved in the regulation of double-strand break repair via nonhomologous end joining, and Jmjd1c, which has histone H3-methyl-lysine-9 demethylase activity. All these genes are speculated to be involved in chromatin organization. In addition, two other genes are worth noting. The first one is Arntl, also known as Bmal1, which is a well-known circadian clock gene. Bmal1 binds E-box enhancer elements upstream of Period (Per1, Per2, Per3) and Cryptochrome (Cry1, Cry2) genes, capable of activating these genes’ transcription. Recently, there has been increasing evidence suggesting the role of biological rhythm in regulating aging (*27-29*). The other gene Apod is a component of high-density lipoprotein (HDL), whose expression is known to increase with age (*30*). The rest of the 23 shared genes are generally related to transmembrane transport, stimulus-response, and regulation of protein folding.

To summarize, the functional enrichment analysis and more detailed DEG analysis suggest that most up-regulated genes are tissue-specific, aimed at improving the specific tissue function. There is considerable overlap between down-regulated genes, enriched for epigenetic modification and transcriptional regulation. Overall, the global transcriptional changes induced by Top2 knock down are beneficial to longevity promotion by acting on several aging hallmarks, as shown in the Circos plot that links DEGs from different tissues to different aging hallmarks (**Fig. 5O; Extended Data Fig. 5G-J**).

### Top2b knock down reprograms the epigenetic landscape and differentially down-regulates highly expressed genes and genes with active promoters

Top2b plays an essential role in regulating DNA replication and transcription by relieving the torsional stress caused by these processes, and its function in regulating transcription is particularly important for post-mitotic cells. To further explore the potential upstream mechanism underlying the longevity effect of Top2b knock down, we analyzed the change of epigenetic landscape and the transcriptional state of DEGs upon Top2b knock down.

We first analyzed various histone markers known to change with age. Compared with the control gLacZ mice, the H3K4me3 (marking active gene promoters and transcription start sites of actively transcribed genes) level in Top2b knockdown mice was significantly down-regulated in kidney, lung, and skeletal muscle. In contract, the inhibitory marks H3K9me3 and H3K27me3 were increased, suggesting that Top2b knockdown leads to global suppression of transcription (**Fig. 6A-D).**

**Figure 6.**
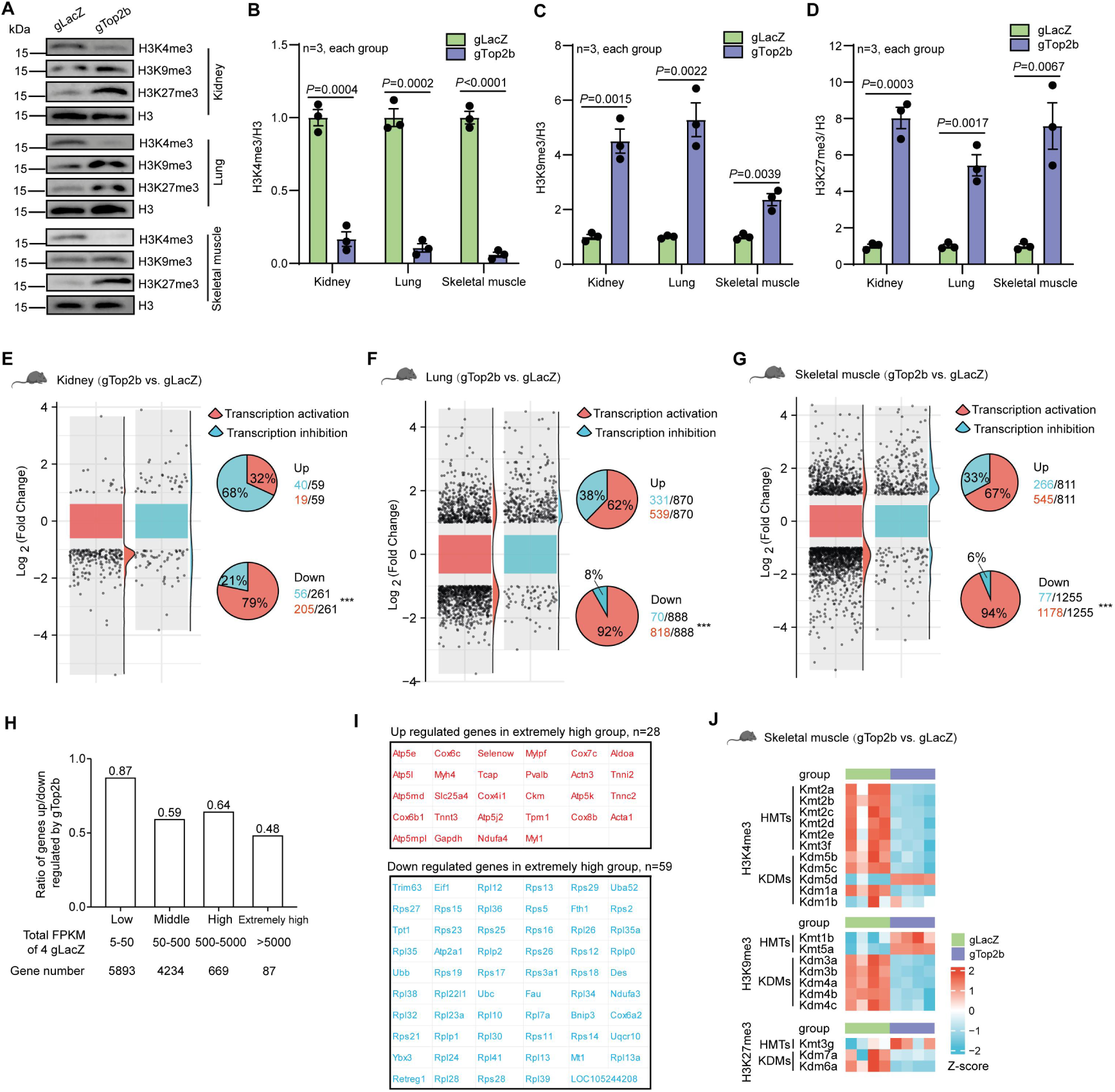
Top2b reduction reprograms the epigenetic landscape and differentially down-regulates genes with active promoters/high abundance. (A-D) Western blot analysis of histone modification markers in mouse kidney, lung, and skeletal muscle. (E-G) The transcriptional state of the promoters of DEGs from mouse kidney, lung, and skeletal muscle. The distribution of the log2(fold change) of active vs. inactive genes (left panel) and percent of active and inactive genes in the up and down regulated DEGs (right panel) were shown. (H) The ratio of genes up-regulated/down-regulated by Top2b knockdown in skeletal muscle decreases with the abundance in gLacZ mice. Genes were grouped into low, middle, high, and extremely high abundance groups according to the total FPKM values in gLacZ mice as indicated. (I) Examples of DEGs with extremely high abundance in muscle. (J) Heatmap depicting the expression levels of genes related to histone methyltransferases (HMTs) and histone lysine demethylases (KDMs) that regulate the trimethylation of histone H3 at lysine residues 4 (H3K4me3), 9 (H3K9me3), and 27 (H3K27me3) in skeletal muscle. Statistical analysis was performed using GraphPad Prism v8.0 software (https://www.graphpad.com). Data were considered statistically significant at *P* < 0.05 calculated by using the Student’s t-test (A and E) or Chi-squared test (E-G). All values are means ± SEM or n (%). The corresponding n values (number of mice) are shown within each sub-plot. *** *P*<0.001.

Interestingly, Top2b knock down seems to elicit a feedback transcriptional response from epigenetic modifiers-histone methyltransferases (HMTs) and demethylases (KDMs) that further reinforce transcriptional suppression. For the active H3K4me3 marker, HMTs are significantly down-regulated while KDMs are up-regulated in muscle (**Fig. 6J**). In contract, for the repressive markers H3K9me3 and H3K27me3, the opposite trend is observed.

We next analyzed whether such transcriptional repression is gene-specific. Numerous studies have indicated that the recognition of Top2b to target genes is not random (*31, 32*). For example, in neurons, Top2b targets are situated within chromatin regions distinguished by H3K4 methylation, and Top2b preferentially binds to promoters, with the occupancy positively correlated with an active transcriptional state (*31*). We thus hypothesized that in aging mice, Top2b knock-down could selectively decrease the expression of those actively transcribed genes, especially those with high abundance.

To address this hypothesis, we utilized the Genome browser (https://genome-asia.ucsc.edu/) to analyze the transcriptional state of the promoter regions of all the differentially expressed genes (DGEs) in mouse kidney, lung, and skeletal muscle (**Supplementary Table 10-12**). In the kidney, the proportion of down-regulated DEGs in the gTop2b group with active transcriptional state in their promoters is 79%, which is significantly higher than the 32% in the up-regulated DGEs (**Fig. 6E, Supplementary Table 10**). Similar results were observed in the lung (**Fig. 6F, Supplementary Table 11**) and skeletal muscle (**Fig. 6G, Supplementary Table 12**). The distribution of log2(fold change) for genes with active transcriptional state shifted significantly towards negative values relative to that of genes with inactive state (**Fig. 6 E-G, left panel**). These results suggest that Top2b knockdown preferentially down-regulates genes with active transcriptional states whose promoters are enriched for active histone marks.

We further analyzed the selectivity of transcriptional suppression at the gene level using the gene expression data from skeletal muscle, which has the strongest transcriptional response based on the total number of DEGs and is relatively homogenous with a smaller number of different cell types. To test whether the transcriptional repression depends on the abundance of gene expression, we group genes into low, middle, high, and extremely high expression groups according to their transcription level in gLacZ mice, and then calculated for each group the ratio between the number of up-regulated and the number of down-regulated genes. We found that while overall there are more down-regulated than up-regulated genes (the ratio < 1), this ratio decreases with gene abundance, indicating that highly expressed genes are more likely to be down-regulated (**Fig. 6H**). Interestingly, for genes in the extremely high group (the total count>5000, **Fig. 6I, Supplementary Table 5**), out of 59 gTop2b down-regulated genes, 41 are ribosomal genes, other 7 are related ubiquitin and autophagy (**Fig. 6I**). Out of 28 gTop2b up-regulated genes, many are involved in muscle development or contraction (**Fig. 6I**). This result from the highly expressed genes is consistent with the global functional annotation of DEGs in skeletal muscle (**Fig. 5M and 5N**).

## Discussion

Top2 is an essential molecular machine that solves DNA topology problems in the cell. The relief of DNA torsional tension by Top2 is required for DNA replication and transcription, thus it is crucial for both proliferative cells and pos-mitotic cells. We found that the reduction of Top2 or Top2b (the mammalian homolog) extends life span across species, improves the health span of mice and worms, and alleviates age-related pathology in various mouse tissues. At the molecular level, Top2b knock down mitigates many hallmarks of aging, including senescence, DNA damage, loss of proteostasis, and de-regulated nutrient sensing. These results suggest Top2 as a novel target for longevity intervention with potentially distinct mechanisms. Our findings also put Top2/Top2b as a fresh example of genes with antagonist pleiotropy in the context of aging – genes important for development and growth but the down-regulation post-development extends health and life span, similar to the classifical insulin/IGF1 and the mTORC pathways (*33*).

We explored the potential mechanisms underlying the prolongevity effect of Top2/Top2b knock down through systematic gene expression profiling of various tissues of mice and analysis of epigenetic markers. Our analyses indicate that selective down-regulation of transcription of genes with active promoter and high abundance is the upstream effect of Top2b knock down. One example is the strong down-regulation of ribosomal genes in muscle, and down-regulation of ribosomal gene expression is a major theme of yeast longevity mutants (*34*). In addition, there is considerable overlap among down-regulated genes in different tissues, including several transcriptional regulators. Intriguingly, such conserved down-regulation can trigger a transcriptional cascade that leads to up-regulation of tissue-specific genes that improve tissue-specific function, such as detoxification in kidney, immune response in lung, and contraction and energy production in muscle. How the conserved down-regulation communicates with tissue-specific transcriptional programs to improve specific tissue function is still unknown and certainly deserves future studies.

Intuitively, Top2b knock down may exert its pro-longevity effect by reducing DNA DSBs, as Top2b is known to co-localize with DNA DSBs which can be caused by failed re-ligation. Although we can not exclude this possibility, our gene expression profiling provided no support for this scenario, with no detectable transcriptional signal for DNA damage response and repair pathways.

Our work suggests that Top2b can be a novel target for longevity intervention. To develop small molecule drugs targeting Top2b, two factors need special attention. One is dosage. Our work indicates that there is an optimal level of knock down of Top2 to best extend life span in both yeast and worm (**Supplementary Table 13**, **Extended Data Fig. 6**), which is not surprising given the essential function of Top2. More importantly, our work suggests that reducing the level of expression of Top2 confers a longevity effect, which may not be achieved by drugs that inhibit the enzyme function at the intermediate steps. A number of topoisomerase inhibitors (such as etoposide, irinotecan, and doxorubicin) have been developed to target cancer; most of these drugs form drug-Top2-DNA cleavage complexes, exacerbating DNA double-strand break (DSB) accumulation (*35-37*). Previous data from yeast suggest such drugs decrease the life span instead of increasing life span (*6*). Thus drugs that specifically target Top2b for degradation such as PROTACs (Proteolysis Targeting Chimeras) are promising candidates (*38*).

## Materials and methods

### Mouse studies ethics statement

Mouse experiments were conducted in compliance with International Guidelines and received approval from the Ethical Committee of the University of Electronic Science and Technology of China, Chengdu, China (Ethics Approval No. 106142024050729362).

### Life span analysis

#### Mice

This study included a total of 38 C57BL/6 mice, among which 30 mice (15 males and 15 females, weighing 20-24g each) were randomly and equally assigned to the gLacZ group and the gTop2b group, with genders distributed almost equally between the two groups. These 30 mice, at 8 months of age, were purchased from SIPEIFU (Beijing) Biotechnology Co., Ltd. [Experimental Animal Production License Number: SCXK (Jing) 2019-0010]. Additionally, we purchased 4 mice aged 2 months and 4 mice aged 10 months from SIPEIFU (Beijing) Biotechnology Co., Ltd. Mice underwent daily health assessments to detect any signs of illness. If a mouse displayed severe distress and was not expected to survive beyond 24 hours, as determined by experienced personnel, it was humanely euthanized. Indicators of severe distress included: (1) inability to consume food or water, (2) pronounced lethargy, characterized by minimal response to external stimuli, (3) significant impairment in balance or movement, (4) rapid weight loss over a week or more, or (5) presence of a severely ulcerated or bleeding tumor. The euthanasia age was documented as the most accurate estimate of the mouse’s natural life span. Deceased mice were noted during daily inspections, and their bodies were preserved for subsequent examinations. For statistical analysis of survivability, the GraphPad Prism v8.0 software (https://www.graphpad.com) was utilized.

#### C. elegans

The N2 (Bristol) wild-type strain was sourced from the Beijing Center for Disease Control and Prevention (Beijing, China) and maintained at a temperature of 20°C. This strain was cultivated on Nematode Growth Medium (NGM) agar plates supplemented with *E. coli* OP50 bacterial lawn. All experiments were performed using L1 synchronized worms. Wild-type worms raised on *E. coli* OP50 underwent bleaching, followed by L1 synchronization in 1X M9 solution for 16–18 hours at 20 °C. Subsequently, the L1 larvae were pelleted at 2500 rpm for 1 minute, the supernatant was removed, and the larvae were transferred to *E. coli* OP50 plates. Upon reaching the late L4 stage, the worms were transferred to RNAi plates. Life span assessment commenced on the 8th day of adulthood and was conducted every other day thereafter. For all RNAi experiments, an empty L4440 vector was used as the control. Statistical analysis of survivability was performed using GraphPad Prism v8.0 software (https://www.graphpad.com).

#### S. cerevisiae

The strains utilized in this investigation, namely BY4741 (*MATα his3Δ1 leu2Δ0 met15Δ0 ura3Δ0*) and BY4742 (*MATα his3Δ1 leu2Δ0 lys2Δ0 ura3Δ0*), are of the *S. cerevisiae* genetic background and have been documented previously. Cultivation of yeast strains was executed in YPD complete medium, comprising 2% glucose, 2% peptone, and 1% yeast extract. Growth of yeast strains occurred in selective minimal media at 30°C with agitation set at 250∼300 rpm. Analysis of replicative life span (RLS) was conducted through a manual microdissection technique, following established procedures. Pre-culturing of strains took place overnight on YPD plates, with all micromanipulation dissections performed at standard laboratory temperature.

### Frailty index (FI) assessment

The FI was evaluated according to established methods. Briefly, each mouse was evaluated for 31 health-related deficits. Each deficit was categorized with a score of 0 (absent), 0.5 (mild), or 1 (severe), as per previous protocols. It’s worth noting that for temperature and weight, scores of 0, 0.25, 0.5, 0.75, or 1 were possible, depending on the deviation from reference values in young adult animals, as outlined in prior literature. The cumulative score was then normalized by the total number of deficits assessed, yielding an FI score ranging from 0 to 1.

### Total RNA extraction and real-time quantitative PCR (RT-qPCR)

Total RNA was extracted from *S.cerevisiae*, *C. elegans,* mouse tissues, and IMR-90 cells using TRIzol reagent (Invitrogen), followed by cDNA synthesis using HiScript III RT SuperMix for qPCR (+gDNA wiper; Vazyme). RT-qPCR was performed following the protocols provided by Applied Biosystems, utilizing a 7500 Fast Real-Time PCR System and the 2 × ChamQ Universal SYBR qPCR Master Mix (Vazyme). Experiments were performed at least three times. The primer sequences are shown in **Supplementary Table 14**. Relative mRNA expression levels were normalized against the respective housekeeping genes: actin (*S.cerevisiae*), actin-1 (*C. elegans*), Gapdh (mouse), or ACTIN (IMR-90) using the 2^-ΔΔCT^ method.

### Western blot analysis

Various whole cell and tissue extracts were obtained with RIPA buffer (Solarbio) supplemented with PMSF and protease inhibitors. Lysates were briefly sonicated and centrifuged at 12000 × g at 4 °C for 20 min. The protein amount was quantified using BCA protein assay (Solarbio) according to the manufacturer’s instructions. A total of 20 µg was separated on 10 or 15% SDS-PAGE gels and then transferred onto a PVDF membrane 20 or 45 μm (Millipore, Inc.). After blocking with 5% Difco Skim Milk (BD Pharmingen) in TBST solution for 1 h, membranes were incubated with specific antibodies overnight at 4 °C. Following antibodies were used: Top2b (1:2000; Catalog# ER65196; HUABIO), Poly (ADP-ribose) polymerase-1 (PARP-1, 1:1000; Catalog# sc-8007; SANTA CRUZ BIOTECHNOLOGY, INC.), TFEB (1:3000; Catalog# 13372-1-AP; Proteintech), mTOR (1:5000; Catalog# ET1608-5; HUABIO), p-mTOR (1:2000; Catalog# HA600094; HUABIO), p70S6K (1:1000; Catalog# HA721354; HUABIO), p-p70S6K (1:1000; Catalog# HA721803; HUABIO), H3K4me3 (1:1000; Catalog# 91264; Proteintech), H3K9me3 (1:5000; Catalog# M1112-3; HUABIO), H3K27me3 (1:1000; Catalog# R26242; ZenBio), H3 (1:5000; Catalog# M1309-1; HUABIO), p16 (1:1000; Catalog# HA721415; HUABIO), p21 (1:1000; Catalog# HA500156; HUABIO), γH2AX (1:1000; Catalog# AF3187; Affinity Biosciences), and β-actin (1:5000; catalog# ET1702-52; HUABIO). After washing, membranes were further incubated for 60 min at room temperature with the secondary antibodies. The signals were detected with Oriscience Supersensitive ECL Kit (Oriscience Biotechnology Co., Ltd). Finally, the membrane was scanned with a Touch Imager System (e-Blot).

### Phylogenetic analysis

The identified amino acid sequence of Top2 in FASTA format was uploaded into MEGA 11 software (https://www.megasoftware.net/), involving sequences from 14 different species. The phylogenetic tree was constructed utilizing the Neighbor-Joining method. Evolutionary distances were computed employing the Poisson correction method, and the nodes of the trees were assessed via bootstrap analysis with 1000 replicates.

### Behavioral tests

#### Rotarod test

Prior to experimentation, mice underwent a comprehensive pre-training regimen spanning 5 days to ensure familiarity with the task. Subsequently, they were carefully positioned on a rotating rod, which underwent acceleration from 4 to 40 rpm over 5 minutes. The precise moments at which the mice lost balance and fell from the drum were meticulously recorded, representing their latency to fall. Each mouse underwent a series of five trials conducted separately, with intervals of no less than 5 minutes between each trial. From these trials, both the average latency to fall and the velocity of falling were calculated with meticulous attention to detail. The Rotarod apparatus employed in these experiments was sourced from Jiangsu Sainence Biotechnology Co., Ltd., China (SA102).

#### Open field test

Four gray arenas (measuring 45 × 45 cm) with elevated walls were employed, maintaining a consistent luminosity of 30 lux throughout the habituation and test phases. Mouse cages were acclimated in the behavior room 60 minutes prior to testing. At the commencement of the experiment, mice were situated within the central region of the arena. Each mouse underwent a 5-minute trial within the arena before being returned to its respective home cage. Subsequently, an overhead camera automatically recorded locomotor activity, with subsequent analysis of behavioral patterns conducted utilizing the SMART Video Tracking System V3.0 (Panlab, Harvard Apparatus).

#### Y-maze test

Before commencing the experiment, mouse cages underwent a 60-minute habituation period within the behavior room. Utilizing a gray Y-maze arena measuring 24.6 cm in length and 7.8 cm in width, consistent luminosity at 30 lux was maintained during both training and test phases. For the Y-maze spontaneous alternation test, each mouse was positioned at one extremity of the same arm and allowed 8 minutes of unrestricted exploration. During the Y-maze novel arm test’s training phase, one arm was obstructed while the mouse was centrally situated within the maze for 10 minutes of initial exploration. Following this, the mouse was returned to its home cage for a 1-hour interval, and the maze was thoroughly cleaned before subsequent trials. In the test phase, the divider was removed, rendering all three arms accessible. The mouse was then placed at the center and granted 5 minutes to navigate the maze freely. Throughout the experiment, the SMART Video Tracking System V3.0 (Panlab, Harvard Apparatus) recorded both the movement trajectories and duration spent within each arm.

#### Elevated zero maze test

The elevated zero maze comprises a circular platform measuring 600 mm in diameter, partitioned into two segments enclosed by walls standing 6 inches high (designated as the ‘shield zone’), and two segments without walls (designated as the ‘open zone’). Mice were introduced at the perimeter of the shielded area, with their movements recorded via an overhead camcorder and subsequently analyzed using the SMART Video Tracking System V3.0 (Panlab, Harvard Apparatus). Each trial lasted for 5 minutes. The percentage of time spent in the open zone served as an index of exploratory behavior and inversely as an indicator of anxiety.

#### Novel object recognition test

Four gray arenas (45 × 45 cm) with elevated walls were utilized, maintaining a constant luminosity of 30 lux throughout both habituation and testing phases. Prior to testing, mouse cages were introduced to the behavioral room for a one-hour habituation period. Subsequently, mice underwent a three-day habituation process: on the first day, co-housed mice were introduced to a single arena for 10 minutes collectively, followed by individual 5-minute sessions for each mouse. This individual exposure continued for days 2 and 3. On the fourth day, mice were individually placed in an arena containing two identical objects positioned equidistant from the walls and each other, allowing for 5 minutes of exploration before being returned to their home cage. After a two-hour interval, mice were reintroduced to the same arena, with one familiar object replaced by a novel one, again given 5 minutes for exploration. Locomotor activity and behavioral patterns were automatically recorded by a camera situated above the arena, and subsequently analyzed using the SMART Video Tracking System V3.0 (Panlab, Harvard Apparatus).

#### Tail suspension test

Mouse cages underwent a 60-minute habituation period in the behavior room preceding the experiment. Subsequently, one mouse per experimental condition was gently secured to the tail using tape and positioned in the tail suspension test apparatus, allowing the mouse to hang upside down. The tail suspension test was conducted for a duration of 6 minutes, during which immobility was precisely quantified using the SMART Video Tracking System V3.0 (Panlab, Harvard Apparatus).

#### Body bending frequency and pharyngeal pumping rate of C. elegans

*C. elegans* specimens were transferred onto an NGM medium devoid of OP50 E. coli. Body bends were quantified as deviations in the movement direction of *C. elegans* (along the X axis), correlated with the movement of the posterior bulb of the pharyngeal ball along the Y axis. Following the stabilization of *C. elegans*, body bending frequency, and pharyngeal pumping rate were documented using stereomicroscopy (Ecoline, Motic) and the Capture 2.1 imaging system (Shenzhen Huaxian Optical Instrument Co., Ltd.).

### Construction of *top2* DAmP strain

The strategy utilized for constructing *top2* DAmP allele followed the previous descriptions (*12*). In brief, the kanamycin-resistance (Kan^R^) cassette was inserted immediately downstream of the open reading frame of *top2* via transformation with a PCR product containing the Kan^R^ cassette, which was flanked at both ends by homologous sequences to the targeted locus.

### RNAi experiments

The *top-2* and HT115 RNAi clones were generously provided by the Beijing Center for Disease Control and Prevention (Beijing, China). All clones underwent thorough verification via DNA sequencing. For RNAi experiments, synchronized populations of animals were cultivated on OP50-seeded NGM plates until the late L4 stage or day 1 of adulthood. Subsequently, they were transferred to RNAi plates (NGM supplemented with 100 ng/μl carbenicillin and 1 mM Isopropyl β-D-1-Thiogalactopyranoside) pre-seeded with bacteria expressing the respective RNAi clone. An empty L4440 vector served as the negative control.

### Cell culture, senescence induction, transfection, adeno-associated virus (AAV) production, and AAV infusion

IMR90 cells were cultured in Dulbecco’s modified eagle medium (DMEM) supplemented with 10% fetal bovine serum (FBS), 1% penicillin/streptomycin, glutamine, sodium pyruvate, non-essential amino acids, and sodium bicarbonate. HEK293FT cells were maintained in DMEM with 10% FBS and 1% penicillin/streptomycin. Regular mycoplasma testing was performed using a LookOut mycoplasma PCR detection kit (Sigma). Senescence induced by oncogenic K-RAS^G12V^ or treatment with etoposide was performed as described.

For transfection of IMR-90 cells, one hour prior to transfection, cells were washed with fresh preheated serum-free DMEM. Then, 1 μg of plasmid was mixed in 50 μl of OptiMEM solution, followed by the addition of 3-5 times the amount of PEI based on DNA quantity. After incubating at room temperature for 15 minutes, the mixture was added dropwise to IMR-90 cell culture dishes. The medium was changed after 4-6 hours. The gRNA sequences targeting LacZ and Top2b were designed from the Cas13design website (https://cas13design.nygenome.org/) and sent to Tsingke Biotechnology Co., Ltd. (Beijing, China) for gRNA synthesis. The gRNA sequences are as follows: gTOP2B-F: aaacTACATCTTCATCATACACCCACA; gTOP2B-R: cttgTGTGGGTGTATGATGAAGATGTA.

Furthermore, the AAV packaging process for intravenous injection into mouse tail veins was conducted as previously described. The gRNA sequences targeting LacZ and Top2b were designed from the Cas13design website (https://cas13design.nygenome.org/) and sent to Tsingke Biotechnology Co., Ltd. (Beijing, China) for gRNA synthesis. The gRNA sequences are as follows: gTop2b-F: aaacTCATGAATAACTTTGAGAGCCAC; gTop2b-R: cttgGTGGCTCTCAAAGTTATTCATGA. Plasmid design involved the selection of the EFS ubiquitous promoter. The AAV-CasRx-Triplex-pregRNA vector was digested with the BSMB1 enzyme and gel purified. The designed gRNAs targeting LacZ and Top2b were annealed at room temperature for 5 minutes, then ligated to the vector with T4 ligase at 16°C for 30 minutes. Escherichia coli were transformed to obtain AAV-CasRx-Triplex-Top2b and AAV-CasRx-Triplex-LacZ plasmids. HEK293FT cells were cultured in DMEM containing 10% FBS; before transfection, the cells were washed with preheated serum-free DMEM. A mixture of 10 μg pHelper, 5 μg DJ, and 5 μg pAAV in 250 μl OptiMEM solution was combined with 100 μg PEI, incubated at room temperature for 15 minutes, and then added dropwise to a 10 cm cell culture dish. The medium was changed after 4-6 hours. After 72 hours, cells were washed with 1× PBS and harvested by centrifugation at 1000 rpm for 3 minutes. AAV virus was extracted from the cells using the AAVpro® Extraction Solution kit (Catalog# 6235, Takara). Fourteen-month-old mice (n=30) were injected via tail vein with AAV-EFS-CasRx-Triplex-Top2b and AAV-EFS-CasRx-Triplex-LacZ viruses, with 15 mice per group, each injected with 100 μl of virus at a titer of 1×10^12^ vg/mL.

### SA-β-Gal assay

SA-β-Gal activity in IMR-90 cells and frozen tissue sections was assessed following the manufacturer’s guidelines (C0602, Beyotime Biotechnology Ltd, Shanghai, China). Analysis and statistical evaluation were conducted using an optical microscope across randomly chosen fields.

### Hematoxylin-eosin (HE) staining

HE staining was conducted using a commercial kit (BH0001, POWERFUL BIOLOGY, Wuhan). Briefly, tissue samples underwent formalin fixation, paraffin embedding, and slide preparation. Deparaffinization was achieved with xylene treatment. Subsequently, the specimens were stained with eosin and hematoxylin to visualize cellular cytoplasm and nuclei, respectively. Following dehydration, sections were analyzed using the Motic Digital Slice Scanning System and Motic DSAssistant software (EASYSCAN, Motic, Xiamen).

### Immunofluorescent staining

IMR-90 cells were cultured on autoclaved coverslips and fixed with 4% paraformaldehyde. Following permeabilization with 0.3% Triton X-100, coverslips were subjected to blocking with a solution containing 10% goat serum and 0.1% Triton X-100 for 1 hour. Primary antibody (γH2AX: Catalog# AF3187; Affinity Biosciences, diluted 1:200) was then applied and incubated overnight at 4 ℃. The following day, coverslips were washed thrice with TBST before incubating with Goat Anti-Rabbit IgG H&L (Alexa Fluor® 594) (Catalog# ab150080, Abcam, China, diluted 1:1000) for 1 hour at room temperature in the dark. Subsequently, coverslips were washed thrice again and immunofluorescence detection was performed using Zeiss LSM800 (Carl Zeiss, Oberkochen, Germany).

### LysoSensor Green and LysoTracker staining in *C. elegans* and IMR-90 cells

For *C. elegans*, worms were immersed in 100 μl of M9 buffer containing 600nM LysoTracker Red DND-99 (LTR, Catalog# 40739ES50, YEASEN, Shanghai, China) and 20μM LysoSensorTM Green DND-189 (LSG, Catalog# 40767ES50, YEASEN, Shanghai, China). Staining was conducted for 1 hour at 20°C in darkness. Subsequently, worms were transferred to NGM plates seeded with fresh OP50 and allowed to recuperate at 20°C for 1 hour in darkness. The fluorescence intensity ratio of LSG to LTR was determined using Zeiss LSM800 (Carl Zeiss, Oberkochen, Germany).

For IMR-90, cells were cultured on autoclaved coverslips and treated with a solution comprising 200 μl of DMEM containing 10% FBS, 60nM LysoTracker Red DND-99 (LTR, Catalog# 40739ES50, YEASEN, Shanghai, China) and 2μM LysoSensorTM Green DND-189 (LSG, Catalog# 40767ES50, YEASEN, Shanghai, China). Staining was conducted in darkness for 1 hour at 37°C. After three subsequent washes, immunofluorescence detection was carried out using Zeiss LSM800 (Carl Zeiss, Oberkochen, Germany).

### Tissue RNA-seq data processing

Transcriptome sequencing was performed on total RNA extracted from the lungs, kidneys, and skeletal muscles of 23-month-old mice (gLacZ group: n=4; gTop2b group: n=4), as well as from *C. elegans* (control group: n=4; *top-2* RNAi group: n=4) at day 8. RNA-seq experiments were conducted utilizing a BGISEQ-500 platform, facilitated by the Beijing Genomic Institution (BGI, China). The sequencing data associated with this study have been deposited in the NCBI Gene Expression Omnibus under the GEO Series accession number GSE278873 (http://www.ncbi.nlm.nih.gov/geo/query/acc.cgi?acc=GSE278873). For gene expression quantification, uniquely mapped reads were normalized to reads per kilobase of exon per million reads mapped, facilitating the calculation of fragments per kilobase of exon per million mapped reads (FPKMs). Differential gene expression analysis identified genes with |log2FC| > 1 and an adjusted *P* value (Benjamini–Hochberg method) < 0.05 as differentially expressed genes (DEGs). The heatmap.2 function from the gplots package version 3.0.1 in R was utilized to generate a graphical representation illustrating Z-score values corresponding to individual genes. DEGs were categorized into upregulated and downregulated genes based on log2FC values for enrichment analysis. Gene Ontology (GO) and Kyoto Encyclopedia of Genes and Genomes (KEGG) pathway analyses were conducted using the R package clusterProfiler. Statistical significance was determined for GO terms and KEGG pathways with a *P* value < 0.05. *P* values were calculated using Fisher’s exact test and adjusted via the Benjamini–Hochberg method. Furthermore, to assess differences in biological process signaling pathways among experimental groups, Gene Set Enrichment Analysis (GSEA) was employed using the R package clusterProfiler. Enrichment analysis result with a *P* value < 0.05 was depicted using the enrichplot package.

### Statistics and reproducibility

The specific sample sizes are provided in the figure legends. Sample size determination did not rely on a predetermined statistical method; however, they align with those documented in prior studies addressing similar topics. Mean values are depicted with standard error of the mean (s.e.m). Statistical significance was evaluated using the Student’s t-test for continuous data and the Chi-squared test for categorical data. Statistical significance was defined as *P* < 0.05. The locally weighted regression (LOESS) function was used to assess the relationship between the mutant gene’s *top2* expression (fold change) and RLS extension in the BY4742 strain. Moreover, with the exception of RNA-seq data, each experiment was independently replicated at least thrice. Detailed descriptions of the statistical methodologies employed, along with precise *P* values, are elucidated in the respective figure legends. No specific techniques were employed for random sample allocation. All data were inclusively analyzed without exclusions. Data collection and analysis were carried out without blinding to experimental conditions. Analysis was conducted using GraphPad Prism v8.0 software (https://www.graphpad.com).

## Acknowledgements

This study was supported by the Sichuan Science and Technology Program (No. 2022ZYD0076, No. 2023YFS0050, No.2024YFHZ0009, No.23NSFSC0349 &No.2023ZYD0066), and by Medico-Engineering Cooperation Funds from the University of Electronic Science and Technology of China and China West Hospital (Grant No. ZYGX2022YGRH018).

## Author’s contributions

M.Z., B.X., J.Y., H.L., and Y.Z. conceived and designed the experiments. M.Z., M.M., FY.L., and LN.L. performed the experiments. M.Z., M.M., LN.L., J.Y., L.Y., Y.X., ZY.W., Y.P., and JS.Z. performed the bioinformatics analysis. M.Z., J.Y., H.L., and Y.Z. wrote the manuscript with input from all authors. All authors read and approved the final manuscript.

